# Approaches to estimating inbreeding coefficients in clinical isolates of Plasmodium falciparum from genomic sequence data

**DOI:** 10.1101/021519

**Authors:** Lucas Amenga-Etego, Ruiqi Li, John D. O’Brien

## Abstract

The advent of whole-genome sequencing has generated increased interest in modeling the structure of strain mixture within clinicial infections of *Plasmodium falciparum* (Pf). The life cycle of the parasite implies that the mixture of multiple strains within an infected individual is related to the out-crossing rate across populations, making methods for measuring this process *in situ* central to understanding the genetic epidemiology of the disease. In this paper, we show how to estimate inbreeding coefficients using genomic data from Pf clinical samples, providing a simple metric for assessing within-sample mixture that connects to an extensive literature in population genetics and conservation ecology. Features of the *P. falciparum* genome mean that some standard methods for inbreeding coefficients and related F-statistics cannot be used directly. Here, we review an initial effort to estimate the inbreeding coefficient within clinical isolates of *P. falciparum* and provide several generalizations using both frequentist and Bayesian approaches. The Bayesian approach connects these estimates to the Balding-Nichols model, a mainstay within genetic epidemiology. We provide simulation results on the performance of the estimators and show their use on ~ 1500 samples from the PF3K data set. We also compare the results to output from a recent mixture model for within-sample strain mixture, showing that inbreeding coefficients provide a strong proxy for the results of these more complex models. We provide the methods described within an open-source `R` package `pfmix`.

## Introduction

While genetic factors play a crucial role in the emergence of drug resistance within (Pf), many aspects of the genetic epidemiology of the parasite remain obscure (Snow et al., 2005; Tibayrenc, 1998). The beginnings of a global perspective on the genetic structure of parasite populations emerged from the analysis of whole-genome sequencing data (WGS) derived from ~ 200 parasite genomes collected directly from clinical patients in six countries on three continents (Manske et al., 2012). This study gave further evidence for the widespread presence of within-isolate strain mixture and significant amounts of variation in its degree across continents. In grappling with the complexity of WGS data, the study departed from standard approaches by attempting to quantify the amount of within-sample variation measure using an inbreeding coefficient, *f_ws_*, a form of F-statistic. Strain mixture is ordinarily assessed via multiplicity of infection (MOI) (Conway et al., 1991; Hill and Babiker, 1995; Hill et al., 1995), using methods for inferring the number of strains from singlenucleotide polymorphisms (SNPs) or other typing technologies applied at a small number of loci. Researchers have subsequently shown how finite mixture models can infer MOI using WGS but the under the heading of complexity of infection (COI) as these model can capture additional mixture features (Galinsky et al., 2015; O’Brien et al., 2015). Still, inbreeding coefficients have a long connection to population genetics and conservation biology and may be of interest to researchers connecting Pf studies to other genetic contexts (Hedrick and Kalinowski, 2000; Weir and Cockerham, 1984). This paper presents a collection of statistical methods for estimating *f_ws_*, details their connection to COI estimates, and confirms the variation in *f_ws_* values across countries using the PF3K data set.

Inbreeding coefficients and the F-statistics from which they derive are measurements of the departure of allelic heterozygosity observed within a population from those expected at Hardy-Weinberg equilibrium (HWE) (Weir and Cockerham, 1984; Nei, 1977). HWE specifies the distribution of alleles assuming panmixia, a population exhibiting perfectly random mating with an absence of mutation, migration, drift, selection or other effects (Wright, 1965). F-statistics calibrate the empirical allele distribution within a subpopulation against those expected under HWE, ranging from a value of one (no mixture) to zero (perfect HWE-type mixture). In the context of comparing the parasites’ genetic diversity within a single infected individual relative to the local geographic population (and absent any geographic structuring of the population, i.e. the Wahlund effect), these statistics effectively become inbreeding coefficients. Specifically, the term *f_ws_* refers to the inbreeding between a subpopulation w and a sample *s*; similarly, *f_iw_* indicates the amount of genetic correlation between a larger popultion *i* and subpopulation *w*. Since we are only concerned here with estimates within a single sample relative to the local population, we will deprecate the paired subscripts and refer to our quantity of interest as *f*, or *f_i_* for a specific sample *i*.

F-statistics have proven to be an effective and extremely popular means for investigating species' population structure from both allelic and genomic data (Weir and Cockerham, 1984; Rousset, 1997; Weir and Hill, 2002). However, standard software tools assume specific ploidy structures incommensurate with WGS data from *P. falciparum* and so cannot be used directly. The critical difference is that, within a human host, Pf exists only in the haploid stage of its life-cycle (Hall et al., 2005). Since short read WGS data cannot yet capture full haplotypes, individual reads cannot be uniquely identified with their strain of origin. Without being able to associate reads to individual *P. falciparum* strains, we cannot see any ‘out-of-the-box’ use of standard *F*-statistics approaches with this new data.

Still, several earlier works connect the framework of F-statistics to Pf within-sample mixture. These concepts – while not under the heading of inbreeding coefficients – undergird much of the seminal work on MOI estimation Hill and Babiker (1995); Hill et al. (1995). More recently, Manske et al. (2012) provides an initial estimator for inbreeding coefficients using WGS based on the slope of a modified regression line between the expected heterozygosity assuming population-level HWE and the observed heterozygosity within a sample. Auburn et al. (2012) explores the connection between this estimator and standard MOI approaches by comparing these estimates with MOI values inferred by genotyping the *msp-1* and *msp-2* genes, showing strong correlation between these values in their sample sets. This paper seeks to clarify this estimator by placing it more firmly within the larger statistical tradition around *F*-statistics.

This paper proceeds as follows. First, we provide an overview of the data and our notation. We present the initial estimator employed by (Manske et al., 2012) for estimating *f_i_* and provide two additional frequentist estimators and detail their connection to classical *F* statistics. We then proceed to describe a Bayesian approach for these statistics that derives from the the Balding-Nichols model. We compare these estimators in a set of simulations. To consider their empirical performance, we examine their correlation in 344 Ghanaian samples and compare the Bayesian estimates to COI estimates. We then present estimates for the PF3K sample set, confirming significant variation in within-sample mixture across countries. We conclude with a brief discussion of the strengths and limitations of our approaches, and possible future directions for modeling within-sample mixture using WGS.

## 1 Data and models

### 1.1 Data and preparation

The data used comes from Release 3.0 of the *Plasmodium falciparum* 3000 Genomes project (PF3K), a publicly available resource of WGS from over 2600 clinical and laboratory Pf samples. An overview of this project, collection protocols, and a full sequencing protocol can be found at the consortial website www.malariagen.net/projects/parasite/pf. For all the samples considered below data come from Illumina HiSeq sequencing applied to clinical Pf samples collected from 14 countries. Starting from the publicly available vcf files, we further excluded samples from Nigeria and Senegal due to sample size and differing sequencing technology, respectively. We further filtered variants by first segregating samples by country and then including only SNPs that exhibited at least 20x coverage at more than 80% of variant positions, and removed SNPs that had at least 20x coverage in all of the remaining samples. This yielded variable number of SNPs within countries, from 1108 in Cambodia to 6596 in Laos. The number of samples within each country ranged from 35 for Laos to 344 in Ghana.

### 1.2 Notation

Within a country, we label the samples *i* = 1,⋯, *N* and the SNPs by *j* = 1,⋯, *M*. At SNP *j* within sample *i*, we observe *r_ij_* reads that agree with the reference, and *n_ij_* reads that are different from the reference. We write *p_ij_* for the allele frequency for reference allele for SNP *j* in sample i and estimate it via the maximum-likelihood estimator (MLE) for proportions: 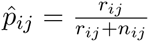. Similarly, we write *p_j_* as population-level reference allele frequency for each SNP and estimate according to the across-sample MLE,

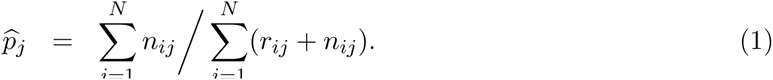

All MLEs are calculated by country. We provide Table 1 as a reference to the reader for notation.

**Table 1.** Notation for parameters used throughout the manuscript.

| Parameter | Description |
| --- | --- |
| $j = 1, \dots, M$ | Index over number of SNPs, $M$ |
| $i = 1, \dots, N$ | Index over number of samples, $N$ |
| $r_{ij}, n_{ij}$ | Reference/non-reference read count data in sample $i$ at variant $j$ |
| $d_{ij} = (r_{ij}, n_{ij})$ | Read count data in sample $i$ at variant $j$ |
| $p_j$ ($\hat{p}_j$) | Population-level non-reference allele frequency for SNP $j$ (estimate) |
| $p_{ij}, \hat{p}_{ij}$ | Within-sample non-reference allele frequency for SNP $j$ in sample $i$, and estimate |
| $f_i$ | Inbreeding coefficient for sample $i$ |
| $H_o(b, i)$ | Observed heterozygosity for sample $i$ in bin $b$. |
| $H_e(b)$ | Expected heterozygosity for bin $b$. |
| $\hat{f}_i^*$ | Estimator of $f_i$ by method $*$. |
| $\mathbf{f}, \mathbf{p}$ | Vector of $f_i$ and $p_j$’s in Bayesian model. |

### 1.3 A previous frequentist estimator for *f_i_*, and two alternatives

In Manske et al. (2012), the authors provide an initial approach to estimating *f_i_*. We refer to this estimator as 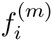 to contrast it with subsequent estimators. For each sample *i*, the estimator first partitions alleles into 10 equally-spaced bins based on their minor allele frequency: (0, 0.05),⋯, (0.45,0.50). Within each bin, *b*, the averaged expected heterozygosity assuming country-level HWE is calculated by

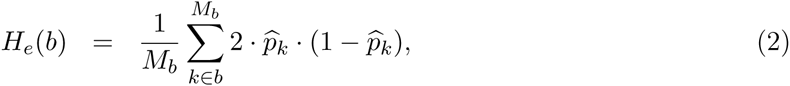

where *M_b_* is the number of SNPs within bin *b*. The averaged observed heterozygosity within each bin and each sample is calculated by

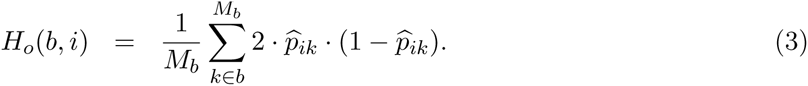

Finally, 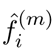 is calculated as 1 – *β* where *β* is the slope found by regressing the *H_o_*(*b, i*) values against 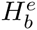 values centered within their respective allele frequency bins and constrained to pass through the origin. This is the **initial estimator**.

The binning procedure, while stabilizing the estimator against influence from an excess of low frequency alleles common within samples, may also introduce estimator bias. We can remove this effect by discarding the binning procedure in favor of directly regressing observed heterozygosity for each SNP against the expected value, still constrained to pass through the origin. This provides a closed-form expression for the **regressed estimator**, 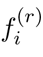, as

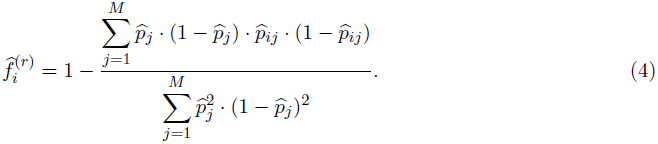

We can also create a similar estimator more transparently derived from the ideas underpinning traditional *F*-statistics in the following way. For a single SNP *j*, suppose *f_i_* to be the fraction of the population-level heterozygosity equal to the difference between the population-level heterozygosity, 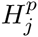 and the sample-level heterozygosity, 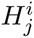 that is,

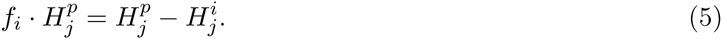

Dividing through by 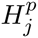 gives an estimate for *f_i_* for the SNP *j*. Averaging across all SNPs and taking the ratio of expectations to be the expectation of the ratios gives the estimator

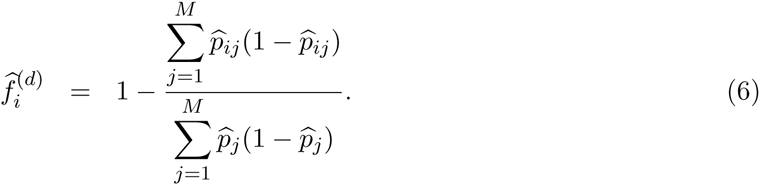

We call this the **direct estimator**, since it contains the critical ratio of the mean observed heterozygosity over the mean expected heterozygosity characteristic of F-statistics.

For each of these estimators, we employ a bootstrap approach to estimate the variance in the estimates for confidence intervals (?Efron and Tibshirani, 1994). The bootstrap works by assuming that the empirically observed distribution – here, the allele frequencies – provides a reasonable approximation to the true empirical distribution. By repeatedly subsampling with replacement from the observed distribution and recalculating the estimator at each iteration, we build up a distribution of estimates from which confidence intervals can be calculated.

### 1.4 Bayesian model framework

We can estimate inbreeding coefficients comparable to the above estimators by employing the Balding-Nichols model, a widely used method for measuring inbreeding in other genetic contexts (Balding, 2003). This approach also has strong similarities to previous work in the context of Pf Hill and Babiker (1995); Hill et al. (1995). In using this model, we make several simplifying assumptions. We treat SNPs as unlinked (i.e. no linkage disequilibrium) and assume that individual parasites within a sample represent a random sample of the surrounding population. We also assume that read counts are sampled identically, independently, and represent an unbiased sample of allele frequency *p_ij_*.

#### 1.4.1 Likelihood and priors

The approach for the **Bayesian estimator** adapts the Balding-Nichols model of allele frequency within inbred subpopulations to the specific context of *P. falciparum* WGS data (Balding, 2003; Balding and Nichols, 1994). In *P. falciparum* the relevant subpopulation is the collection of parasites within a clinical sample. For sample *i* and SNP *j*, Conditional upon an inbreeding coefficient *f_i_* and a population-level allele frequency *p_j_*, the Balding-Nichols model gives the allele frequency *p_ij_* as a Beta distribution,

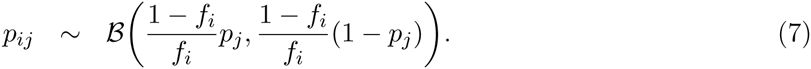

Since the read counts are assumed to be identical and independent, *p_ij_* is drawn from a Beta distribution, and the probability of the data is binomial, we use the conjugacy of these distributions to eliminate the dependence on the unknown *p_ij_* and give a Beta-binomial distribution for the likelihood at a site *j* and position *j*:

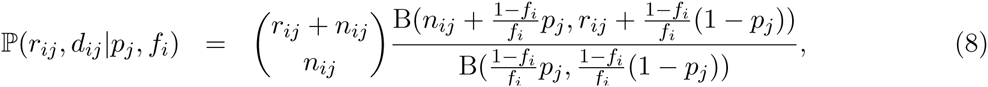

where B(·,·) is the beta function. Since we assume independence by site and by sample, the complete likelihood of the data, *D* conditional upon the inbreeding coefficients for all samples within the population, **f** = (*f*_1_,⋯,*f_N_*) and the allele frequency for all SNPs **p** = (*p*_1_,⋯,*p_M_*) becomes

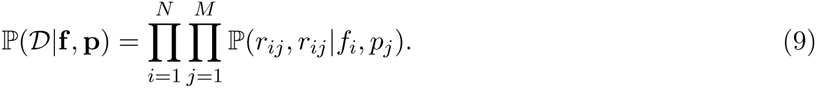

The only prior information we have about the *f_i_* values suggests that high levels of inbreeding are common but not obligatory in west African populations, and we quantitatively interpret this as a uniform prior on each *f_i_* between zero and one. We place a uniform prior distribution on each allele frequency, although we have eliminated rare variants as part of data cleaning described in Section 2.1.

#### 1.4.2 Inference

Since the posterior distribution is not known in closed form, we employ a standard random-walk Metropolis-Hastings Markov chain (MCMC) approach to numerically approximate it (?Gelman et al., 2013). The Metropolis-Hastings algorithm constructs a discrete-time Markov chain over the parameter space in such a way that the posterior distribution of the chain is the stationary distribution of the chain. This requires that at a given iteration in the chain we move from the current parameter state *x* to new parameter state *x*′ with probability *α* given by

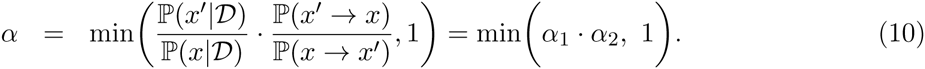

The first ratio is that the posterior probabilities of *x* and *x*′, and we write this as *α*_1_. The second ratio, *α*_2_, gives probability of choosing the current state from the proposed state over the reverse move. Since *α*_1_ constitutes assessment of the likelihood and the prior functions that can be calculated directly from the specifications above, we only describe the calculation for *α*_2_. We denote proposed parameters with an apostrophe.

- **f** - We randomly select *i* and propose *f_i_* from *B* (*α_ci_*, *β_ci_*), leading to 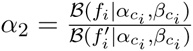.
- **p** - We randomly select *j* and then draw the proposed parameter *p_j_* from the uniform prior, leading to *α*_2_ = 1.
- *α, β* - For both of these parameters, we randomly select individual components and propose new values directly from the prior distribution, leading to 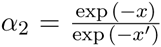 where *x* and *x*′ are the current and proposed state of the relevant component.

The autocorrelation of the log-posterior has minimal lag (Supplementary Figure **??**). As a secondary check, we ran chain for all of the chromosomes individually and compared values with the complete data set. Since we treat SNPs as independent, the performance of the model should be unaffected if the model performs similarly across chromosomes. We find that across all chromosomes performance is nearly identical, with greater than 95% correlation among maximum *a posteriori* (MAP) estimates.

### 1.5 Implementation

All code was implemented in the R computational environment (R Core Team, 2014). The set of scripts implementing each of the estimators, the MCMC algorithm, visualizations, data simulations, and filtered data sets are available at github.com/jacobian1980/pfmix. This repository includes a tutorial and workflow for completing analyses using these approaches. All materials are released under a Creative Commons License.

## 2 Results

### 2.1 Simulations

To compare the qualities of the four estimators, we performed a simualtion study under a range of parameter values to capture how estimator performance may vary with the quality of data collected in the field. We considered the number of SNPs, the number of read counts at each SNP, the degree of skew in the allele frequency distribution, and the amount of inbreeding. For each parameter set, we simulated 100 replicate data sets. The full set of parameters are listed in Table 2.

**Table 2.**
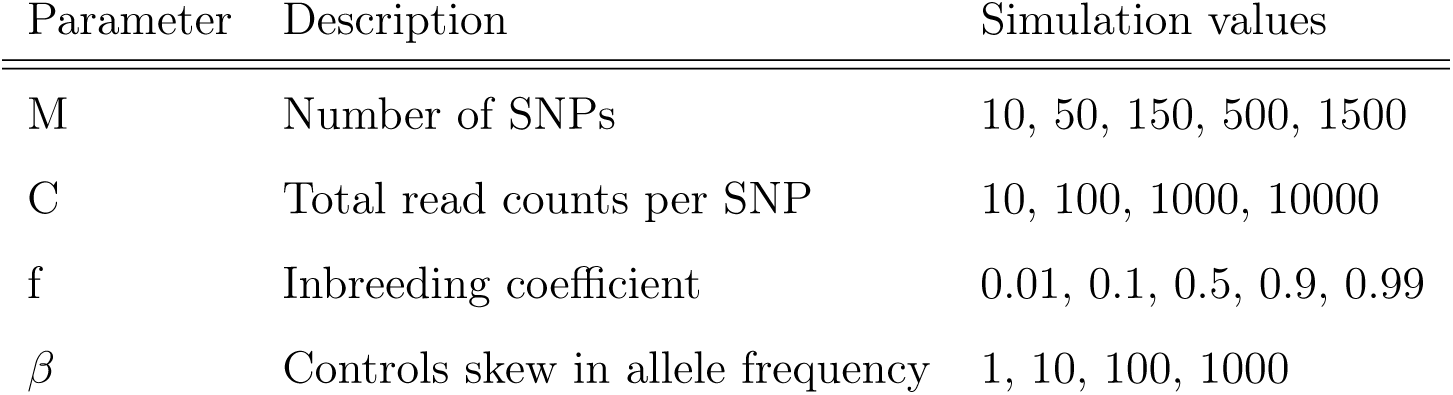
Parameter values for simulated data sets. For each parameter set, 100 replicate data sets were generated.

We simulated data by first fixing the inbreeding coefficient *f* and the allele frequency distribution. We parameterized the allele frequency as a Beta distribution with parametes *α* and *β*. *α* was fixed to one, while *β* was varied according to the simulation to induce differening degrees of skew. As *β* increases, the distribution becomes increasingly right-skewed: when *β* = 1 then 1% of alleles have less than a 0.01 frequency while when *β* = 1000 more than 99% of allele have less than a 0.01 frequency. For a fixed *β* and *f*, we then sample *M* alleles from the distribution and generated observed within-sample allele frequencies according to Equation 7. The read counts were then simulated according to a binomial distirbution with those within-sample allele frequencies.

Figures 1 summarizes the comparison of *f_i_* point estimates made by the initial, regressed, direct, and Bayesian estimators across the simulated data. Performance is recorded as inferred/simulated value. Across all parameter values, the estimators performed similarly, with the Bayesian estimate showing the least bias and highest accuracy. The number of SNPs proves the largest determinant of performance, with 50 SNPs sufficient to ensure reasonable performance in most cases. Very low *f* values (*f* < 0.5) correspond to noticeable bias for the frequentist estimators. The initial estimator is largely robust to large skew in the allele frequency distribution, while the other two estimators are noticeably biased by it at high levels of mixture. We emphasize that the data was simulated under the Balding-Nichols model.

**Figure 1.**
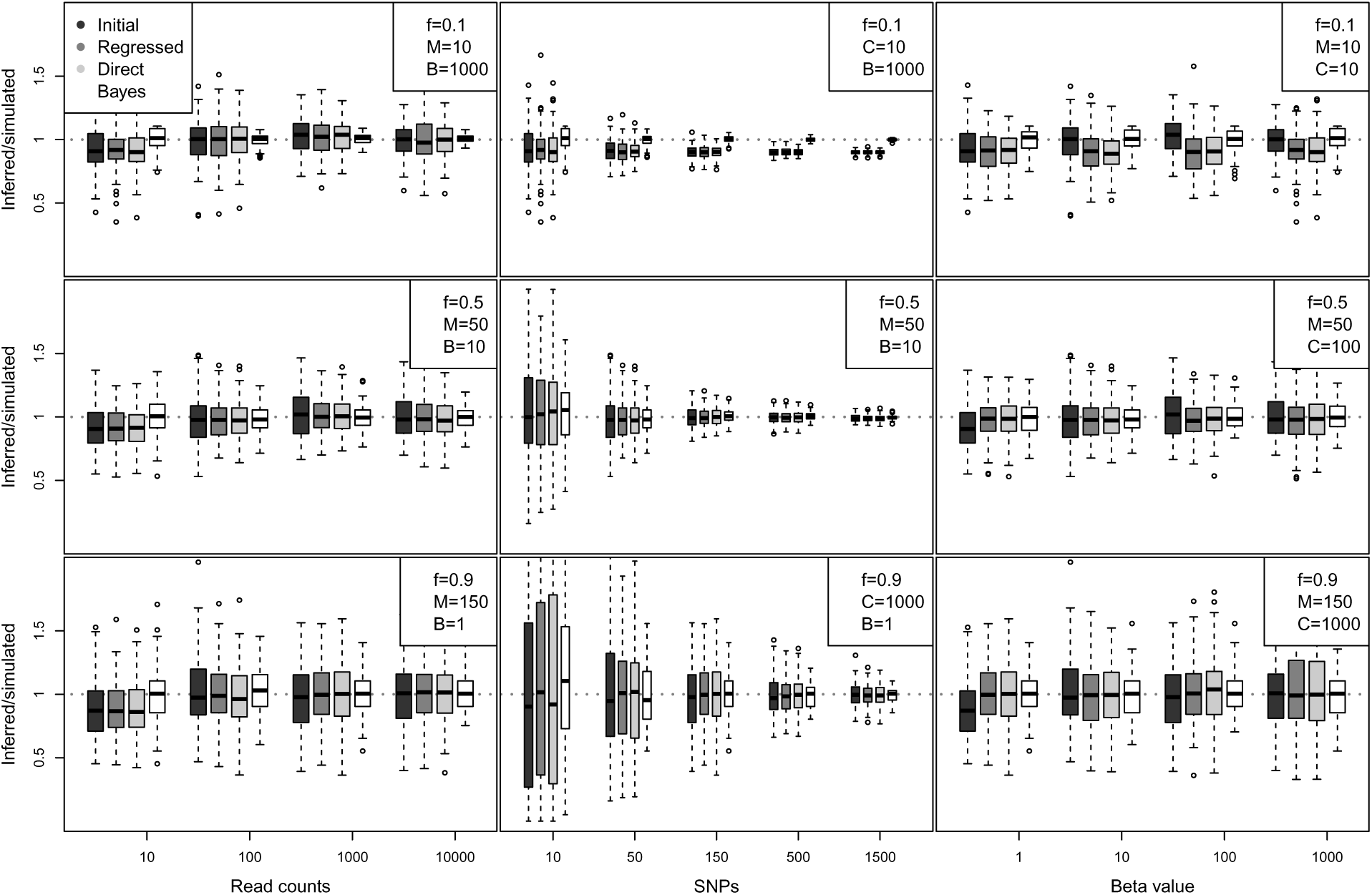
Inferred value over simulated value for each estimator across a range of parameter values. Each boxplot represents 100 replicate data sets with the same parameters.

Figure 2 shows that estimator standard deviation was similar for the three frequentist estimators and markedly smaller in the Bayesian case. For each of the parameter regimes in Figure 1, we performed 100 bootstrap resamplings, even in the Bayesian case. The standard deviation is largely diminished with increasing numbers of SNPs, with read counts and beta values playing little role. We note that bias for the frequentist estimators increases with increasing *f* values.

**Figure 2.**
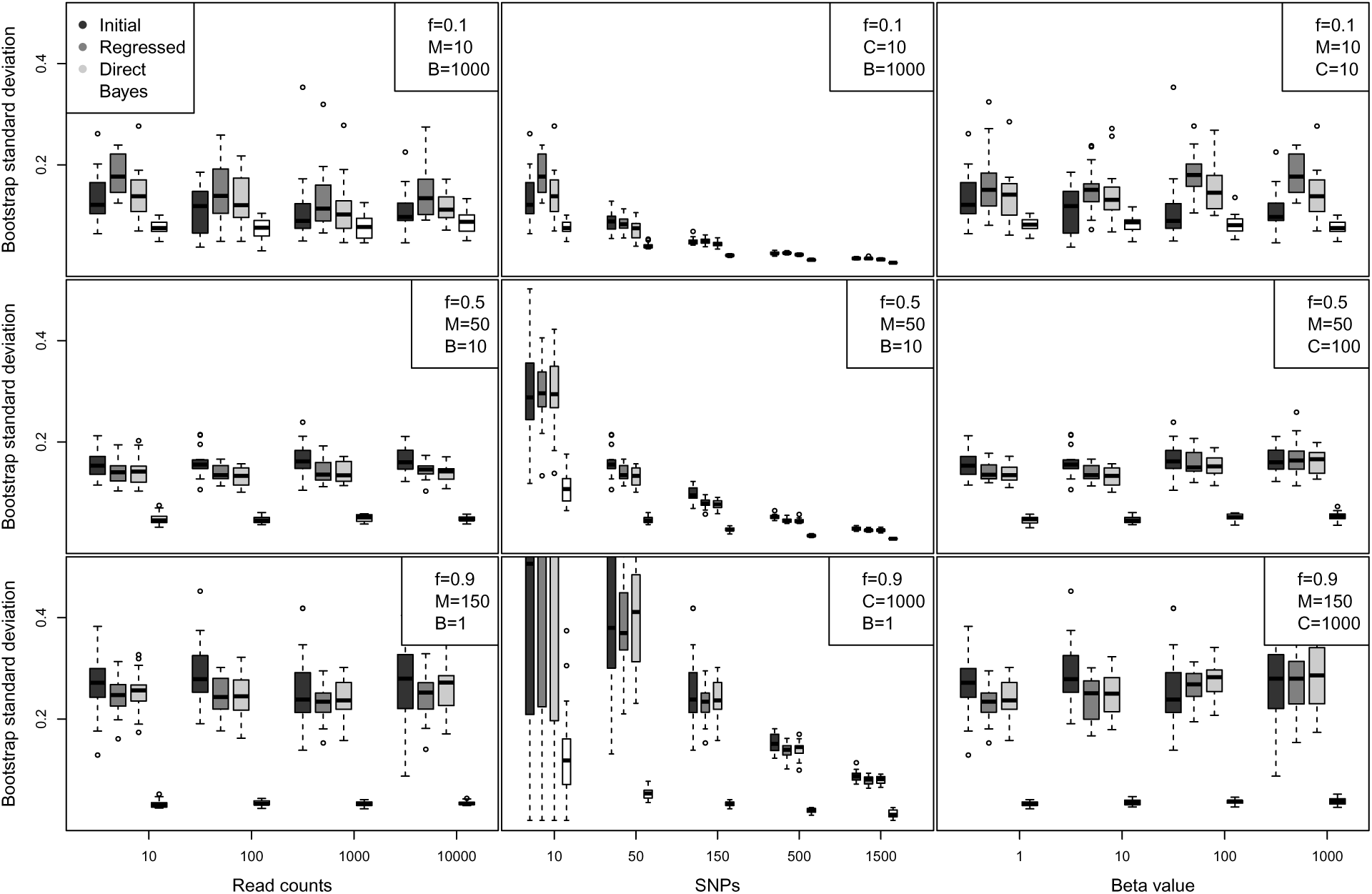
Boostrap standard deviation for each estimator for the same parameter values as Figure 1. Each boxplot represents 50 samples each with 100 replicate data sets.

### 2.2 Comparison in empirical data

Since the underlying Balding-Nichols model within the simulations is likely misspecified relative to empircal data, we examine the performance of the estimators applied to the WGS from 344 Ghanaian samples. The results shown in Figure 3 show very strong correlation between the three frequentist estimators, with correlation better thant 0.95. For the Bayesian estimate, we report the *maximum a posteriori* (MAP) estimate. The Bayesian estimator is still highly correlated (>0.9) with the other estimators but is significiantly more variable in its estimates. Highly mixed and highly unmixed samples (*f* ≈ 0 and *f* ≈ 1) appear to have the most correlation, with moderately mixed samples deviating the most from the other three estimators.

**Figure 3.**
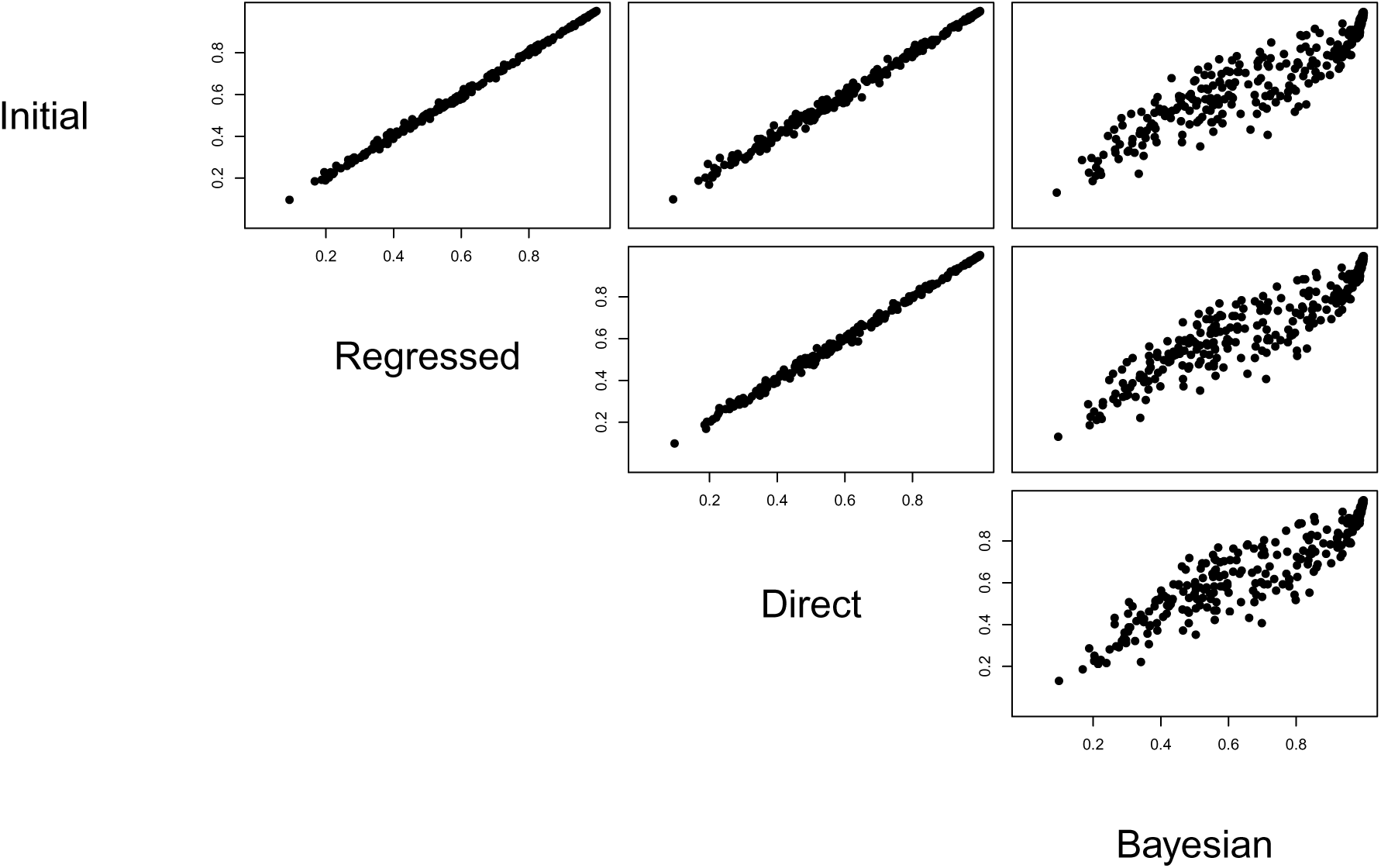
Correlation in inferred *f_i_* value for the four estimators. For the Bayesian case, the MAP value is reported. Each point represents a clinical isolate from Ghana.

### 2.3 Comparison with COI

As noted in the introduction, two recent efforts have extended MOI to WGS and introduced the concept of COI (Galinsky et al., 2015; O’Brien et al., 2015). Both methods use finite mixture models to model the underlying number of strains in the sample. For comparison here, we employ the model of O’Brien *et al*., as it allows for more careful inference of the number of underlying strains and is more robust to errors in the read count data. We apply this model to the 344 Ghanaian samples and for each take the maximum *a posteriori* number of strains as the estimate. Figure 4 shows the correlation between the inferred number of strains and the F-statistic. We calculate a Spearman correlation of 0.83. While complex mixture models may provide a more penetrating understanding of within-sample variance, F-statistics appear to capture much of the same information in a single quantity.

**Figure 4.**
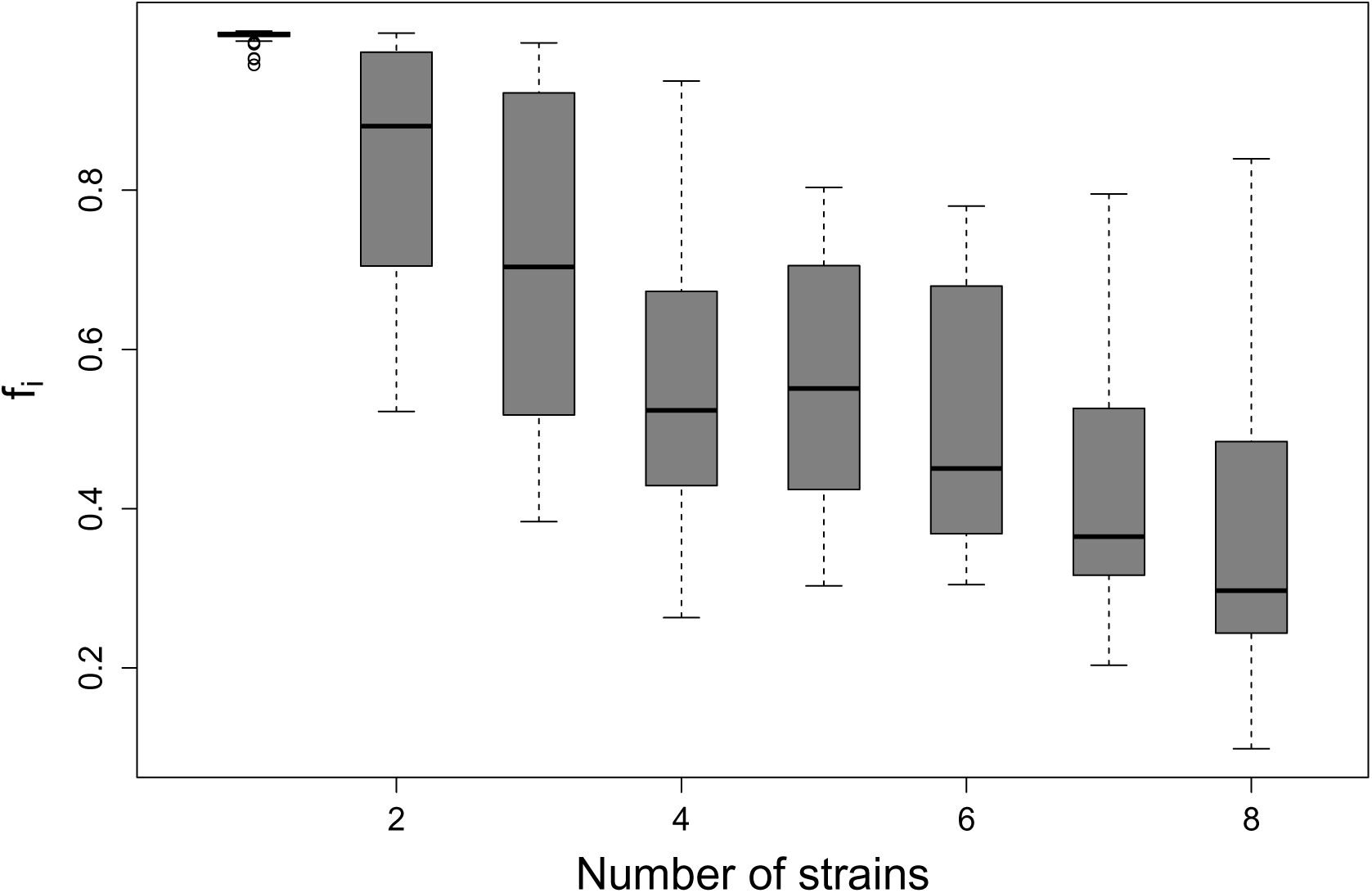
Boxplot of *f_i_* for each sample grouped by number of inferred strains using the mixture model of (O’Brien et al., 2015).

### 2.4 PF3K data set

We grouped the PF3k clinical samples outlined in the Data section by country and used the direct estimator to calculate the inbreeding coefficient for each sample. Figure 5 summarizes the results, showing relatively low *f_i_* values throughout west Africa, with the noticeable exception of The Gambia. This may be due to the geographic segregation of the sampling locations by the Gambia River. The median values of south and southeast Asian countries exhibit distinctly less mixture (higher *f_i_* values) than in West Africa. This is consistent with previous reports of highly variable amounts within-sample mixture across countries (Manske et al., 2012). Interestingly, while the median level of mixture mixture varies significantly across countries, highly mixed samples (*f* < 0.5) and unmixed samples (*f* > 0.95) are present everywhere.

**Figure 5.**
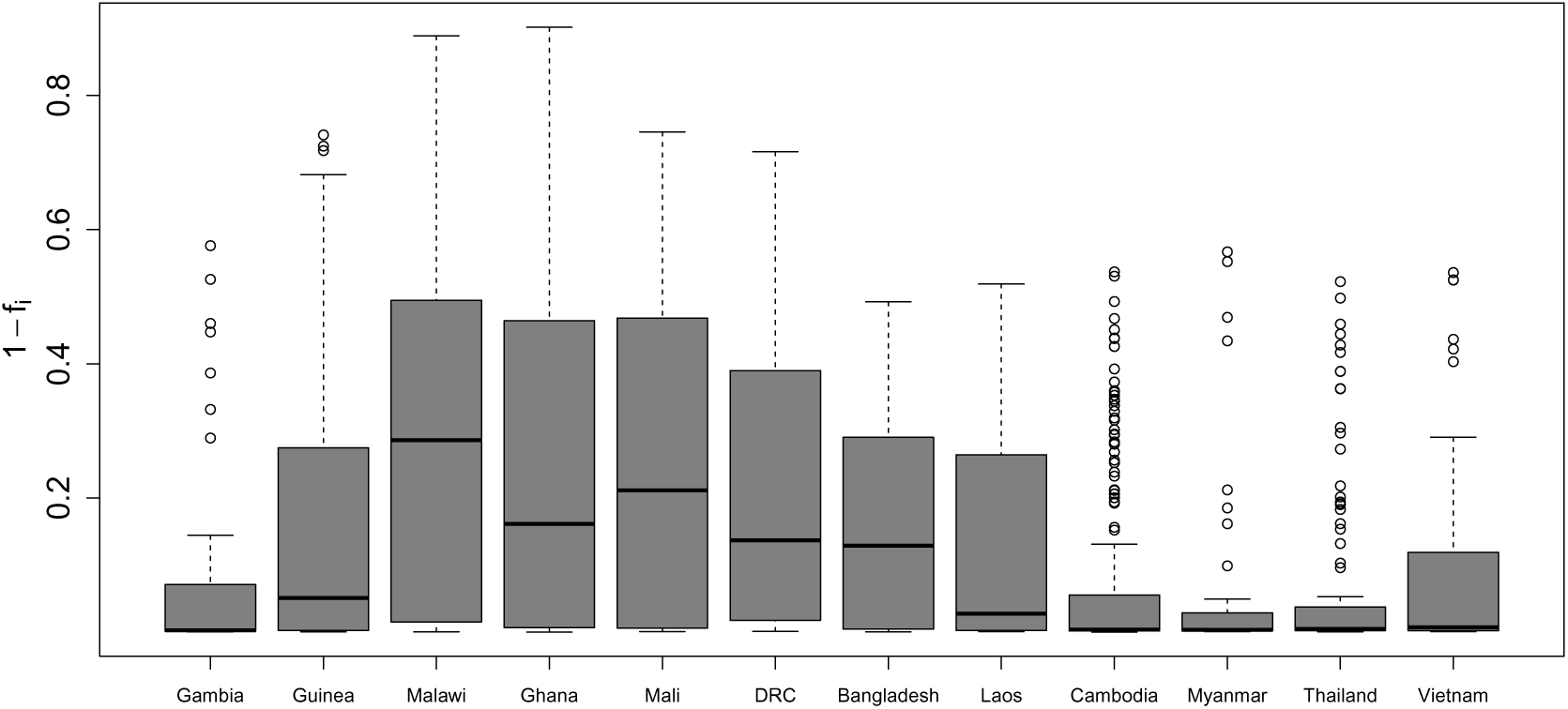
Boxplot of 1 – *f_i_* for each sample grouped by country of origin for 12 countries from the PF3K, arranged from west to east. The more intuitive 1 – *f_i_* is used to emphasize where low and high levels of mixture are prevelant.

## 3 Discussion

This work presents a number of related approaches to inferring inbreeding coefficients for clinical Pf isolates using WGS, using both frequentist and Bayesian approaches. These metrics connect MOI to a broader set of work within population genetics and conservation ecology that may be helpful in control efforts (**??**). We also show that these metrics strongly correlate with more complex mixture models for inferring COI. While we don’t intend for these approaches to supplant these more involved methods for investigating the within-sample mixture, this additional tool can assist researchers in connecting Pf population genetics to a larger literature. To assist other researchers, we also provide the attendant code as an 

~~~
R
~~~

 package, 

~~~
pfmix
~~~

, with additional tutorials and datasets in an open-source context. These are available at the site https://github.com/jacobian1980/pfmix.

The model underlying the inbreeding coefficient makes a number of assumptions about the underlying structure of the read count data and the biological mixing process that may affect our inference. For the read count data, we assume that read counts are unbiased and the SNPs are unlinked. While short read data can be biased in several ways, previous research indicates that mixture proportions calculated by read count ratios are largely unbiased (for instance, see (Manske et al., 2012) supporting information). However, *P. falciparum* exhibits significant linkage disequilibrium on scales significantly larger than the average distance between neighboring SNPs in our data. We do not expect this violation to bias our estimates as this absence of independence occurs (roughly) evenly across the genome. However, inference from a small region of the genome will likely exhibit bias.

A perhaps more troublesome assumption is embedded in the underlying structure of the *F*-statistic. An *F*-statistic measures the departure of the observed number of heterozygotes relative to those expected under Hardy-Weinberg equilibrium. In the context of mixed *P. falciparum* infections, the equilibrium assumptions – random mating, no selection, large population size, genetic isolation – are likely each violated at some level. For example, the mixture within a sample may be the result of a small number of founding individuals or be strongly selected by the human immune system. Without a more general approach to understanding the mixing process, we cannot anticipate the robustness of our estimates to this sort of misspecification. However, we do find that the PF3K samples from Cambodia that possess quite significant population structure still exhibit strong correlation between *f_i_* and the inferred number of strains.

As genomic data enables more elaborate statistical models for mixed infections and a broader understand of Pf genetic epidemiology, it will still be useful for field researchers to connect their work with population genetics and ecology through simple metrics. Inbreeding coefficients, which have a history going back to the beginnings of modern genetics, connect to a number of population genetic quantities such as effective population size and genetic drift (Hedrick and Kalinowski, 2000; Lande and Barrowclough, 1987; Nei and Tajima, 1981) and may serve to complement traditional MOI values and newer models to this end. This work meets this need by providing a basis for calculating these quantities and a suite of open-source tools for researchers.

## Author's contributions

JO’B designed and implemented the study and wrote the manuscript. RL provided visualization of the data and model results. LA-E collected the data, contributed to the study design, and edited the manuscript.

## Acknowledgements

We are grateful for many helpful discussions with Jason Wendler. This publication uses data from the MalariaGEN Plasmodium falciparum Community Project: www.malariagen.net/projects/parasite/pf.

## Supplementary figures

**Figure S1.**
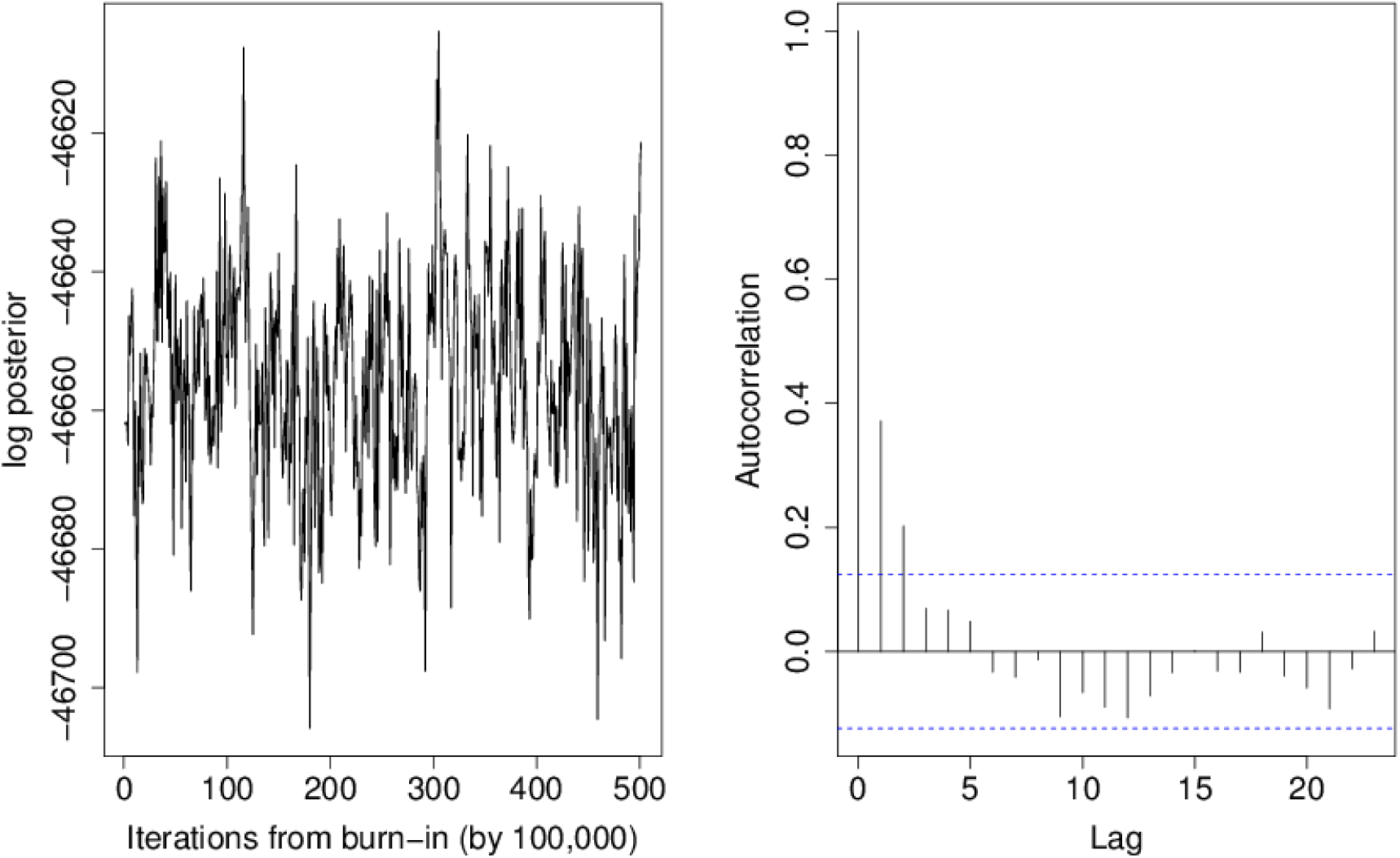
Log-likelihood trace (left) and autocorrelation for MCMC chain for the Bayesian model.

## References

Auburn, S., Campino, S., Miotto, O., Djimde, A. A., Zongo, I., Manske, M., Maslen, G., Mangano, V., Alcock, D., MacInnis, B., et al. (2012). Characterization of within-host *Plasmodium falciparum* diversity using next-generation sequence data. PloS one, 7(2):e32891.

Balding, D. J. (2003). Likelihood-based inference for genetic correlation coefficients. Theoretical population biology, 63(3):221–230.

Balding, D. J. and Nichols, R. A. (1994). Dna profile match probability calculation: how to allow for population stratification, relatedness, database selection and single bands. Forensic Science International, 64(2):125–140.

Conway, D., Greenwood, B., and McBride, J. (1991). The epidemiology of multiple-clone plasmodium falciparum infections in gambian patients. Parasitology, 103(Pt 1):1–6.

Efron, B. and Tibshirani, R. J. (1994). An introduction to the bootstrap. CRC press.

Galinsky, K., Valim, C., Salmier, A., de Thoisy, B., Musset, L., Legrand, E., Faust, A., Baniecki, M. L., Ndiaye, D., Daniels, R. F., et al. (2015). Coil: a methodology for evaluating malarial complexity of infection using likelihood from single nucleotide polymorphism data. Malaria journal, 14(1):4.

Gelman, A., Carlin, J. B., Stern, H. S., Dunson, D. B., Vehtari, A., and Rubin, D. B. (2013). Bayesian data analysis. CRC press.

Hall, N., Karras, M., Raine, J. D., Carlton, J. M., Kooij, T. W., Berriman, M., Florens, L., Janssen, C. S., Pain, A., Christophides, G. K., et al. (2005). A comprehensive survey of the plasmodium life cycle by genomic, transcriptomic, and proteomic analyses. Science, 307(5706):82–86.

Hedrick, P. W. and Kalinowski, S. T. (2000). Inbreeding depression in conservation biology. Annual review of ecology and systematics, pages 139–162.

Hill, W. G. and Babiker, H. A. (1995). Estimation of numbers of malaria clones in blood samples. Proceedings of the Royal Society of London. Series B: Biological Sciences, 262(1365):249–257.

Hill, W. G., Babiker, H. A., Ranford-Cartwright, L. C., and Walliker, D. (1995). Estimation of inbreeding coefficients from genotypic data on multiple alleles, and application to estimation of clonality in malaria parasites. Genetical research, 65(01):53–61.

Lande, R. and Barrowclough, G. F. (1987). Effective population size, genetic variation, and their use in population management. Viable populations for conservation, pages 87–123.

Manske, M., Miotto, O., Campino, S., Auburn, S., Almagro-Garcia, J., Maslen, G., O’Brien, J., and Kwiatkowski, D. (2012). Analysis of *Plasmodium falciparum* diversity in natural infections by deep sequencing. Nature, AOP.

Nei, M. (1977). F-statistics and analysis of gene diversity in subdivided populations. Annals of human genetics, 41(2):225–233.

Nei, M. and Tajima, F. (1981). Genetic drift and estimation of effective population size. Genetics, 98(3):625–640.

O’Brien, J. D., Iqbal, Z., and Amenga-Etego, L. (2015). An integrative statistical model for inferring strain admixture within clinical plasmodium falciparum isolates. arXiv preprint arXiv:1505.08171.

R Core Team (2014). R: A Language and Environment for Statistical Computing. R Foundation for Statistical Computing, Vienna, Austria.

Rousset, F. (1997). Genetic differentiation and estimation of gene flow from f-statistics under isolation by distance. Genetics, 145(4):1219–1228.

Snow, R. W., Guerra, C. A., Noor, A. M., Myint, H. Y., and Hay, S. I. (2005). The global distribution of clinical episodes of *Plasmodium falciparum* malaria. Nature, 434(7030):214–217.

Tibayrenc, M. (1998). Genetic epidemiology of parasitic protozoa and other infectious agents: the need for an integrated approach. International journal for parasitology, 28(1):85–104.

Weir, B. S. and Cockerham, C. C. (1984). Estimating F-statistics for the analysis of population structure. evolution, pages 1358–1370.

Weir, B. S. and Hill, W. (2002). Estimating f-statistics. Annual Review of Genetics, 36(1):721–750.

Wright, S. (1965). The interpretation of population structure by f-statistics with special regard to systems of mating. Evolution, pages 395–420.

